# The interplay of spontaneous pupil-size fluctuations and EEG power in near-threshold detection

**DOI:** 10.1101/2024.05.13.593918

**Authors:** Veera Ruuskanen, C. Nico Boehler, Sebastiaan Mathôt

## Abstract

Detection of near-threshold stimuli depends on the properties of the stimulus and the state of the observer. In visual detection tasks, improved sensitivity is associated with larger pre-stimulus pupil size. However, it is still unclear whether this association is due to optical effects (more light entering the eye), correlations with arousal, correlations with cortical excitability (as reflected in alpha power), or a mix of these. To better understand this, we investigated the relative contributions of pupil size and power in the alpha, beta, and theta frequency bands on near-threshold detection. We found that larger pre-stimulus pupil size is associated with improved sensitivity and a more liberal criterion (the likelihood of reporting that a stimulus was detected). Importantly, the relationship between pupil size and sensitivity was not mediated by any of the measured neural variables; the relationship with criterion however was mediated by power in the beta band (12-30 Hz). Pupil size was positively correlated with power in the alpha and beta bands. Additionally, we show that pre-stimulus theta suppression (4 - 8 Hz) is associated with improved sensitivity and a more liberal criterion. Taken together, our results show an independent effect of pupil size on detection sensitivity that is not driven by cortical excitability, but may be driven by optical effects, arousal, or a mix of both.

---

The interplay of spontaneous pupil-size fluctuations and EEG power in near-threshold detection

Whether a stimulus is perceived or not depends on the external properties of the stimulus and the internal state of the observer. Internal factors are especially relevant at the threshold of detection, where the same stimulus is sometimes perceived and sometimes not. In this context improved sensitivity is associated with larger pupils (Eberhardt et al., 2022; Mathôt & Ivanov, 2019) and changes in the frequency spectrum of the electroencephalogram (EEG), especially in pre-stimulus alpha power (Ergenoglu et al., 2004). These effects are related, but do not necessarily reflect the same underlying mechanism. Enhanced visual sensitivity can result from optical effects (pupil dilation increases visual sensitivity by allowing more light into the eye) as well as neural effects (enhancement of cortical excitability increases visual sensitivity by changing how visual input is processed). Here, we attempt to dissociate the two.

The state of the brain affects sensory perception. One aspect of ‘brain-state’ is cortical excitability, indexed by oscillations in the alpha frequency band (8 - 13 Hz) in the EEG. Lower power in the alpha band (alpha suppression) has been suggested to indicate higher excitability (Lange et al., 2013; Romei et al., 2008). Both pre-stimulus alpha power (measured at occipital electrodes) (Benwell et al., 2017, 2022; Ergenoglu et al., 2004; Limbach & Corballis, 2016; Melcón et al., 2024; Samaha et al., 2017) and alpha phase (Busch et al., 2009; Mathewson et al., 2009; van Dijk et al., 2008) have been associated with near-threshold detection. Concerning alpha specifically, lower pre-stimulus levels are associated with better performance (Ergenoglu et al., 2004; van Dijk et al., 2008). There is some debate as to whether alpha-suppression enhances visual sensitivity itself (Ergenoglu et al., 2004; Romei et al., 2010; van Dijk et al., 2008) or simply modulates conscious awareness of a presented stimulus (Benwell et al., 2017, 2022; Samaha et al., 2017). But regardless of the exact mechanism, there is clear evidence that neural oscillations related to cortical excitability are linked to detection performance.

Another—largely neglected—factor that influences detection performance is pupil size. Studies manipulating pupil size through changes in background brightness show improved detection of faint stimuli in the visual periphery when pupils are larger (Eberhardt et al., 2022; Mathôt & Ivanov, 2019). Similarly (although to a lesser extent) smaller pupils are related to improved discrimination performance (Campbell, 1957; Campbell & Gregory, 1960; Mathôt & Ivanov, 2019; Woodhouse, 1975). Larger pupils benefit detection by increasing the amount of light that enters the retina, leading to a better signal-to-noise ratio. On the other hand, smaller pupils benefit discrimination by focusing incoming light on the fovea, where visual acuity is highest (Mathôt, 2020). Interestingly, in an experiment involving both detection and discrimination, average pupil size was found to be larger in the detection condition, despite no other obvious differences (i.e., in difficulty or background brightness) between conditions (Mathôt & Ivanov, 2019). This suggests that the pupil is able to flexibly adjust to a size that is optimal to the demands of the task at hand (for further discussion see (Vilotijević & Mathôt, 2023).

The pupil does not only respond to changes in visual input but also to internal factors such as cognitive activity. Pupil-size changes have been observed in a variety of tasks and contexts, such as problem solving, working memory, decision making, and attention (Alnæs et al., 2014; Hess & Polt, 1960, 1964; Kahneman & Beatty, 1966; Keene et al., 2022). In general, increases in task difficulty or processing load lead to pupil dilation irrespective of the specific cognitive processes involved, and larger pupils are often associated with better performance (for reviews see Beatty, 1982; Laeng et al., 2012; Loewenfeld, 1958; Mathôt, 2018; Vilotijević & Mathôt, 2023). These types of pupil responses are suggested to be mainly driven by changes in arousal (Bradley et al., 2008; Gilzenrat et al., 2010; Unsworth & Robison, 2018). This follows from the link between pupil size and activity in brain regions associated with arousal regulation, most notably the locus coeruleus (LC) (Joshi et al., 2016; Joshi & Gold, 2020; Murphy et al., 2014). In addition to arousal regulation via norepinephrine release, the LC is implicated in other high-level cognitive functions, such as attention and behavioural control (Aston-Jones et al., 1999; Aston-Jones & Cohen, 2005). Due to the link between pupil size and LC-activity, pupil size is often used as a marker of arousal.

Furthermore, fluctuations in pupil size are correlated with alpha power both during task performance and at rest. For instance, in an auditory oddball task, increased alpha prior to stimulus presentation is associated with decreased stimulus-evoked pupil dilation (Hong et al., 2014). Similarly, in a visual detection task, alpha power and pupil size show a positive correlation within the pre-stimulus interval (Pilipenko & Samaha, 2024). During rest, or inactive wakefulness, studies show associations between alpha power and various aspects of pupil dynamics. Montefusco-Siegmund et al. (2022) have linked alpha power with high-frequency pupil diameter fluctuations and the rate of change in pupil size, while Ceh et al. (2020) found correlations with both the average and variance of pupil diameter. Specifically, they report a positive relationship between alpha power and average pupil diameter, contrasted by a negative relationship with the variance in pupil diameter. Despite these nuances, the general trend indicates a positive correlation between pupil size and alpha power.

Taken together, previous research has shown that both alpha power and pupil size are associated with detection sensitivity but in different ways: alpha power and detection sensitivity show a negative correlation, while pupil size and detection sensitivity show a positive correlation. Furthermore, as outlined above, alpha power and pupil size are positively correlated. This pattern of results—a triangle of seemingly inconsistent correlations—suggests that alpha power and pupil size affect detection performance through different routes. Indeed, in a study on visual detection and confidence across multiple stimulus contrast levels, alpha power and pupil size were shown to have dissociable behavioural effects, whereby alpha modulated detection criterion (i.e. the likelihood to report the presence of a target regardless of whether a target was actually presence) and pupil size modulated detection sensitivity (Pilipenko & Samaha, 2024).

More specifically, lower pre-stimulus alpha power was associated with a lower criterion across all contrast levels, and larger pre-stimulus pupil size was associated with improved sensitivity, especially at higher contrast levels. To our knowledge, no other studies have explicitly investigated the relative effects of alpha power and pupil size on detection. Yet, in the experiment by Pilipenko and Samaha (2024) described above, as in many other studies on pupil-linked effects, pupil size is assumed to function simply as a marker of arousal. Following this assumption, the large-pupil advantage observed in near-threshold detection would be attributed to arousal variation. However, this account overlooks the potential optical effect of pupil size itself. As discussed above, pupil size controls how light falls on the retina, leading to changes in visual acuity, which is likely to play a role in determining detection performance.

Furthermore, both spontaneous and experimentally induced changes in pupil size have been shown to affect neural signatures of visual processing. These include the amplitude of the C1-component, associated with processing in the primary visual cortex (Bombeke et al., 2016), activity patterns in the beta-frequency range (Mathôt et al., 2023), and the strength of steady-state visual evoked potentials (SSVEPs) (Suzuki et al., 2019). These findings suggest that pupil size changes optimise vision at the earliest stages of visual processing. Still, research explicitly investigating pupil-related effects on perception is relatively scarce.

Here, we aim to further investigate the relative contributions of pupil size and neural activity on near-threshold detection, with a focus on the often overlooked effect of the pupil. In terms of neural activity, in addition to alpha-power we will also analyse activity patterns in the beta and theta frequency bands. We expect to find that both larger pre-stimulus pupil size and lower pre-stimulus alpha power are associated with better detection performance. To investigate the relative contributions of pupil size and power in the different frequency bands we will conduct a mediation analysis. However, due to the exploratory nature of this study, we do not formulate specific hypotheses on the nature or direction of possible mediation effects.

## Methods

### Open-practices statement

Experimental materials, raw data, and analysis scripts are available on the open-science framework (https://osf.io/9p563/). The study was not pre-registered.

### Participants

19 participants with normal or corrected-to-normal vision participated in the experiment. Sample size was not predetermined and is arguably moderate compared to other studies, but reflects the number of participants that VR was able to test during a research visit to the lab of NB. The two participants who did not have normal vision wore soft contact lenses (not glasses) during the recording. After preprocessing, 3 participants were excluded from the analysis due to poor data quality (10% or more of epochs marked bad and high amounts of noise identified already during recording). Data exclusion was conducted based on visual inspection, and decided before any further analyses had been conducted. The mean age in the final sample (N = 16) was 24 years, and there were 5 males and 11 females.

All participants provided written informed consent prior to participation. The experiment was approved by the ethics committee of the psychology department at the University of Gent (study code: SEP 2021-215). Participants received a €25 monetary compensation for participating in the full experiment (€15/hour during task performance and €10/hour during EEG preparation).

### Detection task

Participants completed a near-threshold visual detection task consisting of reporting the presence of a faint peripherally presented stimulus. The experiment and stimuli were created with and controlled by OpenSesame (version 3.3.9, Lentiform Loewenfeld) (Mathôt et al., 2012).

All stimuli were presented on a grey (RGB = 128, 128, 128) background with a luminance of 18.5 cd/m^2^. A circular grey fixation dot (RGB = 89, 89, 89) with a size of 0.3° of visual angle (20 px) was maintained in the centre of the screen throughout the experiment (Figure 1).

The to-be-detected target stimulus was a white luminance patch (RGB = 255, 255, 255) with a Gaussian envelope with a standard deviation of 0.46° (30 px) (Figure 1). The luminance of the stimulus at full contrast was 36.4 cd/m^2^. However, the contrast and spatial frequency of the stimulus were adjusted with a Quest adaptive staircase procedure to keep overall accuracy fixed at 75%. Consequently, stimulus-luminance during the task was lower. Possible values ranged between 5% and 25% for contrast, and 0.1° and 0.2° of visual angle for spatial frequency. Stimulus location was determined by drawing a random angle between 0° and 360° with a fixed eccentricity of 7.57° of visual angle (500 px).

Each trial started with a variable target-onset time drawn from a flat distribution between 1 and 2 seconds. Next, on half of the trials the target stimulus was flashed for 50 ms. After target onset each trial lasted for an additional 1.5 seconds. Therefore, total trial length varied between 2.5 and 3.5 seconds. Since the fixation dot was maintained on the screen throughout the experiment, it was not clear to the participants when an individual trial started or ended. Participants were instructed to press the spacebar on the keyboard whenever they detected the target. Upon keypress the fixation dot changed to a dot with a Gaussian envelope for 10 ms to indicate to the participant that a response was recorded.

The task consisted of a practice phase and experimental phase. There were 12 trials in the practice phase and 1040 trials in the experimental phase. The trials of the experimental phase were divided into 16 blocks, separated by self-paced breaks. During 4 of the blocks (block numbers 1, 5, 9, and 13), the Quest adaptive staircase procedure was applied to vary the properties of target stimulus as described above to keep overall accuracy fixed at 75%. These blocks consisted of 50 trials each, and were not included in the analysis. The remaining 12 blocks consisted of 70 trials each, yielding a total of 840 experimental trials that were included in the analysis. Participants received feedback on their average accuracy at the end of each block. One block lasted approximately 3 to 4 minutes.

The experiment was presented on a 27-inch (68.58 cm) Benq LED monitor (model number: XL2411Z) running at a refresh rate of 100 Hz and 1920 x 1080 resolution. The viewing distance was approximately 100 cm for all participants.

**Figure 1:**
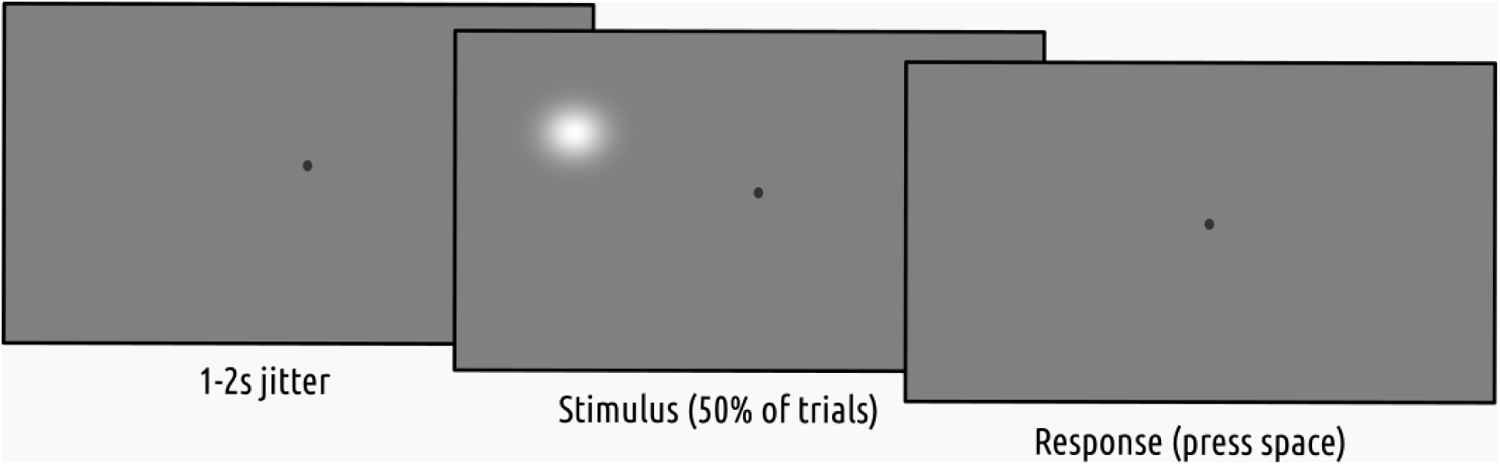
Detection task. This illustration is not to scale. Stimulus contrast as shown is higher than in the actual experiment for illustration purposes.

### EEG acquisition

The EEG signal was acquired from 64 Ag/AgCI scalp electrodes, positioned according to the international 10-20 system, as well as three external electrodes, placed on the mastoids and under the left eye. The signal was recorded at 512 Hz, using a BioSemi ActiveTwo amplifier with its built-in CMS-DRL recording logic, and the associated software (BioSemi B.V.). No online referencing or filtering was done (other than the standard anti-aliasing filter related to the sampling rate). EEG was measured continuously from the beginning to the end of the experimental task.

### Pupil size measurement

Pupil size was measured with an Eyelink 1000 eye tracker (SR Research Ltd., 2022) with a 35 mm lens. An initial 9-point calibration procedure was performed at the beginning of the experiment, and a recalibration was performed halfway through. A drift correction was performed at the beginning of each block. Pupil size was measured from the beginning to the end of each trial. Participants rested their head on a chinrest throughout the experiment, keeping the distance to the eye tracker fixed at 60 cm.

### Procedure

Upon arrival in the lab, participants received information about the study in both verbal and written form, after which they signed the informed consent form. After the informed consent procedure the EEG cap was placed on the participants’ head, filled with gel, and optimised until all electrode offsets were low (< 10 kΩ) and stable, and the eye tracker was calibrated.

Next, participants completed the detection task. Specific task instructions were given verbally before the informed consent procedure, and again on the screen at the beginning of the task. Participants were tested in a Faraday cage with constant, dim illumination (13 lux). The light-source in the room was a small table lamp behind the screen, which fully covered it from direct view, yielding a diffuse illumination.

The whole experimental session, including informed consent, preparation, task performance, and debriefing, took approximately 2 hours per participant.

### Preprocessing

#### EEG

EEG preprocessing was conducted in the MATLAB (version 9.11 R2021b) (*MATLAB*, 2021) toolbox EEGLAB (version 14.1.1) (Delorme & Makeig, 2004). The data was visually inspected for exceptionally large artefacts, which were removed if found. Then, the data was re-referenced to the mastoids, and a 0.1 Hz high-pass filter was applied. Independent component analysis (ICA) was run for each participant, and topographic maps of the components were created. The signal was reconstructed without blink related components (identified through visual inspection from the ICA topographic maps), and the components were then rejected following a visual inspection of the reconstructed signal. Possible bad channels (noted during recording) were excluded from the ICA. They were subsequently interpolated and reincorporated to the data.

Next, data was exported from MATLAB and imported into eeg_eyetracking_parser, a Python library for analysis of combined EEG and eye tracking (Mathôt et al., 2023), which in turns relies on MNE (Gramfort et al., 2013), Autoreject (Jas et al., 2017), EyeLinkParser (Mathôt and Vilotijević, 2022), and DataMatrix. Data was downsampled to 100 Hz. When extracting epochs, the Autoreject algorithm (Jas et al., 2017) was applied to detect and interpolate bad epochs and channels that had not been identified in the previous preprocessing steps. Autoreject relies on cross-validation to estimate the optimal peak-to-peak threshold in the data (for details, see (Jas et al., 2017).

### Pupil size

Missing or invalid data was interpolated using cubic-spline interpolation if possible, using linear interpolation if cubic-spline interpolation was not possible (when the segment of missing data was too close to the start or end of a trial), and removed if interpolation was impossible altogether (when data was missing from the start and/ or until the end of a trial) or if the period of missing data was longer than 500 ms and thus unlikely to reflect a blink. Next, pupil size was z-scored for each participant separately, across all blocks.

### Analysis

#### Time-Frequency analysis

To characterise the power dynamics of the pre-stimulus interval, we conducted a time-frequency analysis on the EEG signal, focusing on posterior and parietal channels (O1, O2, Oz, POz, Pz, P3, P4, P7, P8). We computed the power spectrum for single trials across a frequency range of 4 Hz to 30 Hz with a fast-fourier transform using Morlet wavelets, as implemented in MNE (function tfr_morlet) (Gramfort et al., 2013). The epoch used in the time-frequency analysis spanned from 2.5 s before to 0.5 s after stimulus onset. For further analysis, power was normalised by z-scoring for each frequency band and participant separately, and the epoch was cropped (to avoid filtering edge artefacts) to span 1 s before stimulus presentation.

#### Mediation analysis

To examine the relative contributions of pupil size and neural activity on detection performance, we used structural equation modelling (SEM) to conduct a mediation analysis using the lavaan (Rosseel, 2012) package in R (R Core Team, 2021). The aim was to test whether power in the alpha, beta, and theta frequency bands mediates the relationship between pupil size and detection performance. We specified a model that included paths from pupil size to each potential mediator, from each mediator to the dependent variable (in two separate analyses: accuracy and proportion of target-present responses), and a direct path from pupil size to the dependent variable. Indirect effects were computed as the path coefficients from pupil size to each mediator and from the mediator to the dependent variable, while the direct effect is represented by the path coefficient from pupil size to the dependent variable. The model was fitted separately for each participant. We then conducted both traditional and Bayesian (Dienes, 2014) one-sample t-tests to evaluate the significance of the parameter (slope) estimates against a null hypothesis of no effect. All models were constructed using the average power and pupil size in the 1-second interval before stimulus onset.

## Results

### Descriptives: behavioural performance

The descriptive statistics on behavioural performance measures including average accuracy (i.e. hits and correct rejections), reaction time (RT), and d-prime (d’) are presented in Table 1. D’ is a measure of perceptual sensitivity, defined as the difference between z-transformed hit and false alarm rates. Due to some participants having a false alarm rate of 0, leading to infinite d’ values, we applied a so-called log linear correction (Hautus, 1995). This approach involves first adding 0.5 to both the number of hits and false alarms and then adding 1 to the total number of both signal (i.e., stimulus present) trials and noise (i.e., stimulus absent) trials, prior to computing d’.

**Table 1.** Descriptive statistics of behavioural performance.

| Statistic | M | SD | Min | Max | N |
| --- | --- | --- | --- | --- | --- |
| Accuracy | 72.07 % | 6.62 | 62.08 % | 84.66% | 16 |
| RT (ms) | 569.14 | 97.93 | 357.49 | 683.95 | 16 |
| $d'$ | 1.91 | 0.5 | 1.15 | 2.54 | 16 |

### Descriptives: pupil size

The distribution of z-scored pupil sizes recorded during the experiment is depicted in figure 2a. There was a significant positive relationship between average pupil size (across participants) and trial number (*r* = 0.47, *p* < 0.001), indicating that average pupil size increased slightly over the course of the experiment (figure 2b). Average pupil size was also larger after breaks (indicated by dashed lines in figure 2b) and then decreased over the course of a block. However, despite variations throughout the block, we included all experimental trials in our analysis to avoid excluding genuine fluctuations in pupil size.

**Figure 2:**
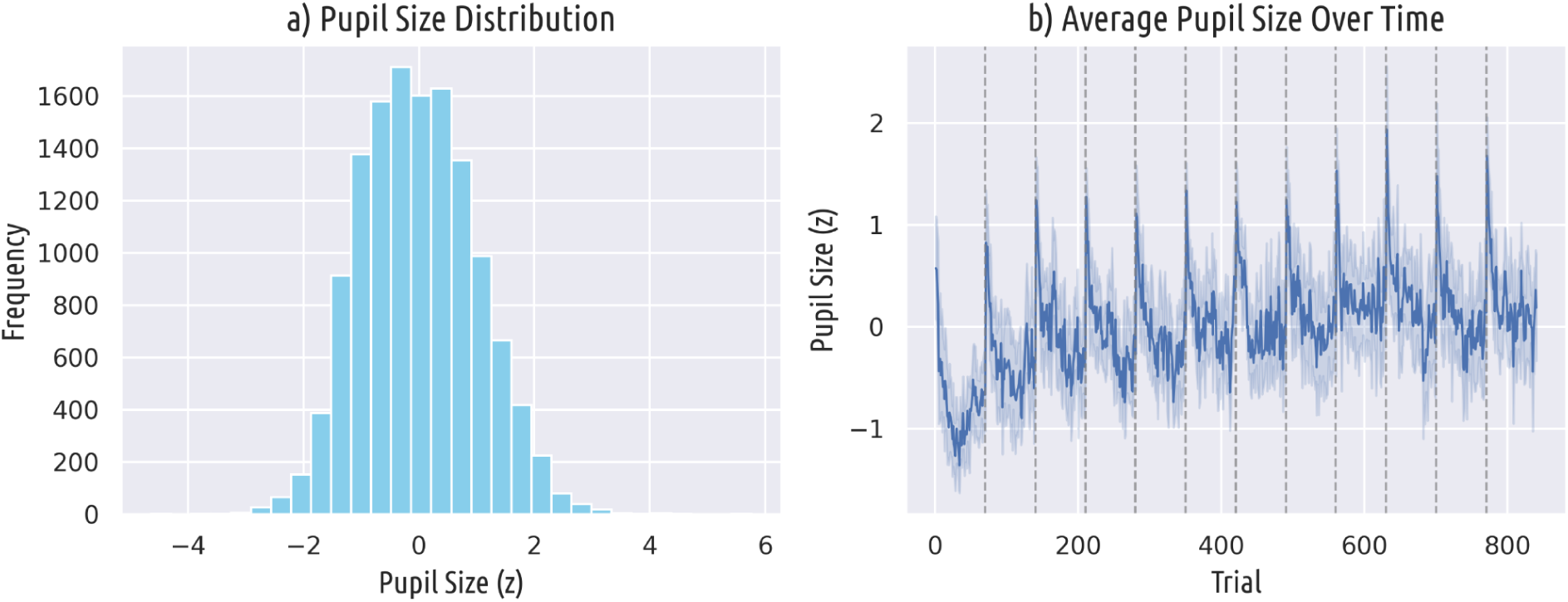
Pupil size descriptives. A) Distribution of recorded pupil sizes. B) Average pupil size over time. Dashed vertical lines represent block-breaks.

**Figure 3:**
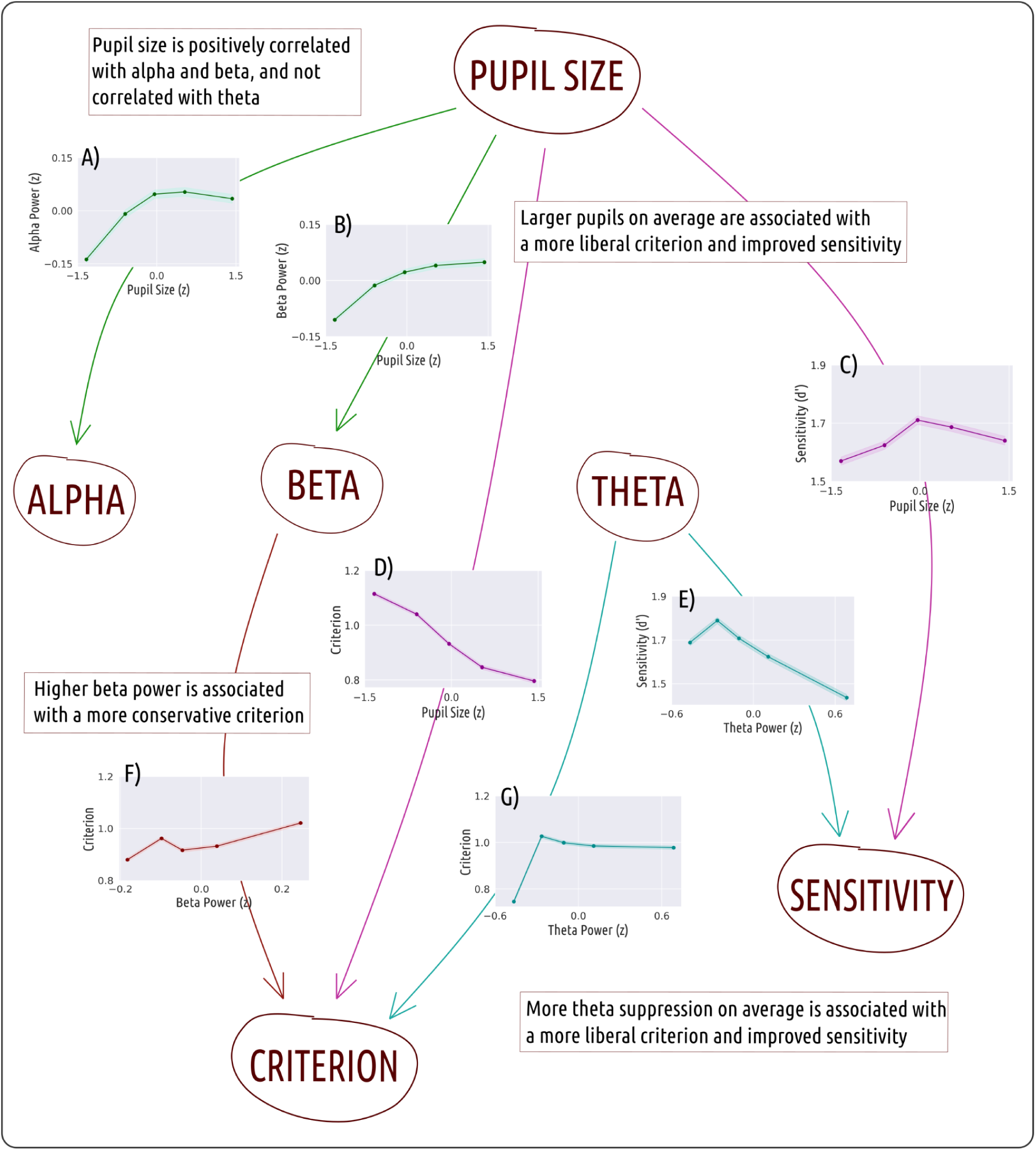
Mediation analysis results. Arrows represent significant effects. Arrows and figures pertaining to the same result are colour-coded. A) Pupil size is positively correlated with power in the alpha frequency band. B) Pupil size is positively correlated with power in the beta frequency band. C) Medium-to-large pupil sizes are associated with improved sensitivity (d’). D) Larger pupils are associated with a more liberal criterion. E) Theta suppression is associated with improved sensitivity (d’). F) An increase in beta power is associated with a more conservative criterion. G) Theta suppression is associated with a more liberal criterion.

### Larger pupils are associated with improved sensitivity and a lower criterion

Our mediation analysis considered two single-trial dependent measures: accuracy (whether the response was correct) and target-present response (whether the target was reported as present regardless of whether it actually was). These dependent measures roughly map onto respectively response sensitivity (d’) and criterion, such that a high d’ indicates high accuracy and a low criterion indicates a high number of target-present responses. We will use d’ and criterion for visualisation (Fig 3); however, these cannot be calculated for single trials and thus not used as dependent measures in our mediation analysis.

Larger pupils were associated with increased accuracy on single trials (*b* = 0.02, *t(15)* = 2.86, *p* = 0.012) and consequently, higher sensitivity, visualised in Figure 3c as a relationship between binned pupil size and d’. Larger pupils were also associated with an increased proportion of target-present responses (*b* = 0.03, *t(15)* = 3.98, *p* = 0.001), visualised in Figure 3d as a relationship between binned pupil size and criterion.

These results were corroborated by the Bayesian analysis, showing moderate to strong evidence for the alternative hypotheses (i.e., path coefficient differs from zero). The direct pathway from pupil size to accuracy showed a Bayes Factor of 4.71, while the direct pathway from pupil size to response showed a Bayes Factor of 32.79.

### Pupil size is positively correlated with alpha and beta power, but not theta power

Larger pupils were associated with increased power in the alpha (Fig 3a) and beta (Fig 3b) bands, but not the theta band. Direct effects from pupil size to alpha power (*b* = 0.05, *t(15)* = 2.39, *p* = 0.030) and from pupil size to beta power (*b* = 0.05, *t(15)* = 4.03, *p* = 0.001) were significantly different from zero, but the direct effect from pupil size to theta power was not (*b* = -0.00, *t(15)* = -0.32, *p* = 0.750).

The Bayesian analysis provided moderate to strong evidence for the alternative hypothesis (i.e., path coefficient differs from zero) for the relationships between pupil size and alpha, and pupil size and beta. The path coefficient from pupil size to alpha power showed a Bayes Factor of 2.22 and the path coefficient from pupil size to beta power showed a Bayes Factor of 35.54. For the relationship between pupil size and theta power on the other hand, the Bayes Factor in favour of the alternative hypothesis was 0.27, indicating weak support.

### Theta suppression is associated with improved sensitivity and a lower criterion

A decrease in power in the theta frequency band was associated with increased accuracy on single trials (*b* = -0.03, *t(15)* = -3.56, *p* = 0.002) and consequently, higher sensitivity visualised in Figure 3e as a relationship between binned power and d’. Theta suppression was also associated with an increased proportion of target-present responses (*b* = -0.02, *t(15)* = -2.93, *p* = 0.010) visualised in Figure 3g as a relationship between binned power and criterion.

Bayesian analyses showed moderate to strong support for the alternative hypotheses (i.e., coefficient differs from zero). The path coefficient from theta to accuracy showed a Bayes Factor of 15.51 and the path from theta to response showed a Bayes Factor of 5.29, both suggesting the presence of significant effects.

### The relationships between pupil size and sensitivity are not mediated by power in the EEG

The indirect effects in our mediation model with accuracy as a dependent variable did not significantly differ from zero. This was indicated both by traditional one sample t-tests and Bayesian t-tests. Specifically, the Bayesian analysis provided moderate evidence for the null hypothesis across all three indirect pathways. The path coefficient from pupil size to accuracy via beta power showed a Bayes Factor of 3.03 in favour of the null; similarly, the path through alpha power showed a Bayes Factor of 2.90, and the pathway through theta power had a Bayes Factor of 3.91, all suggesting the absence of significant indirect effects. These results indicate that the relationship between pupil size and sensitivity was not mediated by power in any of the EEG frequency bands.

### The relationship between pupil size and criterion is mediated by power in the beta band

Our mediation model with response as the dependent variable showed a slightly different pattern of results. Specifically, there was a significant direct effect of beta power on response (Fig 3f; visualised as the relationship between beta and criterion; *b* = -0.03, *t(15)* = -2.35, *p* = 0.032) and a significant indirect effect of pupil size on response via beta (*b* = -0.001, *t(15)* = -2.16, *p* = 0.047). Bayesian analyses provided moderate to anecdotal evidence for the alternative hypotheses (i.e., coefficient differs from zero). The direct path from beta power to response showed a Bayes Factor of 2.10 and the indirect path from pupil size to response via beta showed a Bayes Factor of 1.56, suggesting the presence of significant effects.

However, the indirect effects of pupil size on response via alpha and theta did not significantly differ from zero. Specifically, the Bayesian analysis provided anecdotal to moderate evidence for the null hypothesis across both indirect pathways. The path coefficient from pupil size to accuracy via alpha power showed a Bayes Factor of 1.57 in favour of the null; similarly, the path through theta power showed a Bayes Factor of 3.67, both suggesting the absence of significant indirect effects.

## Discussion

Here, we have investigated the effects of pupil size and neural activity on visual detection performance. We observed that both theta suppression and increased pupil size are associated with improved detection sensitivity. Importantly, these effects are distinct, meaning that the effect of pupil size on detection sensitivity is not mediated by power in the theta band (nor either of the other two frequency bands investigated here). Further, both pupil size and theta suppression are associated with a more liberal criterion and the relationship between pupil size and criterion is mediated by power in the beta frequency band. Finally, we have shown a positive correlation between pupil size and power in the alpha and beta bands, replicating previous research (Mathôt et al., 2023; Montefusco-Siegmund et al., 2022; Pilipenko & Samaha, 2024). Our findings suggest a complex interplay between pupil size, arousal, and visual detection, revealing a network of relationships that is less straightforward than previously thought.

Contrary to expectations based on previous research (Ergenoglu et al., 2004; Montefusco-Siegmund et al., 2022; Pilipenko & Samaha, 2024; Samaha et al., 2017), we found no significant relationship between alpha power and either sensitivity or criterion. Some authors have suggested that pre-stimulus alpha phase is more relevant in determining stimulus detectability than alpha power is (Busch et al., 2009; Mathewson et al., 2009; van Dijk et al., 2008), which may explain our findings. However, we did observe a pattern in the theta range similar to what is typically seen in alpha: theta suppression was associated with higher sensitivity and a more liberal criterion. While the literature on the role of theta power in visual perception is sparse, there is evidence suggesting that theta phase can influence stimulus detection (Busch et al., 2009; Somer et al., 2020), and like alpha power (Lange et al., 2014), theta power has been linked to perceptual decision-making in contexts of illusory perception (Mathes et al., 2014). This suggests that mechanisms related to visual perception may be comparable across these two frequency ranges.

The key finding of our study is that larger pupils benefit detection independently of the changes observed in the EEG power spectrum. Pupil-size related effects on visual detection are often overlooked, and thus, for the remainder of this section we will focus on discussing these results. The improvement in visual sensitivity that results from increased pupil size can be explained in two ways. Firstly, larger pupils allow more light into the eye, thus decreasing the signal-to-noise ratio and improving sensitivity (Mathôt, 2020), which we call the optical effect. Secondly, larger pupils reflect variations in arousal (or cortical excitability), which affects visual sensitivity by changing the way the stimulus is processed, which we call the arousal-effect. Dissociating these effects is not trivial and is further complicated by the lack of consensus on what ‘arousal’ is and which measures reflect it in which ways (Eysenck, 1976; Robbins, 1997; Ross & Van Bockstaele, 2021).

Another complicating factor in dissociating these effects is that the relationship between pupil size and arousal is more nuanced than previously understood. Traditionally viewed as a direct marker of arousal due to its association with locus coeruleus (LC) activity (Joshi et al., 2016; Murphy et al., 2014), recent findings suggest this connection is more variable (Megemont et al., 2022). Research done in mice by Megemont et al. (2022) reveals that only large and infrequent pupil dilations correspond directly with spiking activity in the LC, challenging the notion of pupil size as a consistent readout of LC activity. Additionally, variability in this relationship might be influenced by behavioural factors, highlighting inconsistencies in how pupil dilation reflects arousal states. Furthermore, another study examining arousal-related neural activity and pupil size in mice found that dynamic changes in pupil size, rather than absolute size, align more closely with neural activity changes associated with different arousal states in the dorsal lateral geniculate nucleus (dLGN). More specifically, tonic spiking (associated with wakefulness) occurred more during pupil dilation, while burst spiking (associated with low-arousal states) occurred more during pupil contraction (Crombie et al., 2024). These findings underscore the complexity in linking pupil size directly to arousal-related neural activity.

Somewhat similarly, while changes in the EEG frequency spectrum have been used as markers of cortical arousal (Danos et al., 2001; Foucher et al., 2004) or cortical excitability (Pilipenko & Samaha, 2024), different studies use different terms (e.g., arousal, excitability, activation) and define different EEG correlates. Some treating arousal and cortical excitability as interchangeable terms, while others do not, is one reflection of the lack of a clear definition of arousal as a unitary concept (Eysenck, 1976). However, insofar as amplitude differences in the lower frequencies can be considered to reflect arousal-variation (Foucher et al., 2004) we have shown that these do not mediate the relationship between pupil size and detection sensitivity. This supports the assumption that a part of the large-pupil advantage observed in near-threshold detection is due to an optical effect driven by pupil dilation itself.

Additional evidence for the optical effect comes from behavioural experiments measuring pupil size in detection and discrimination tasks, as well as pupil-size measurements obtained in experiments on auditory near-threshold detection. As previously mentioned, in a study involving both detection and discrimination, average pupil size was found to be larger in the detection condition, despite no obvious differences in task difficulty (Mathôt & Ivanov, 2019). The lack of a difference in difficulty implies that there should be no systematic differences in applied mental effort or arousal. While it is theoretically possible that visual detection tasks are inherently more arousing than discrimination tasks, in a standard lab-setting this seems unlikely. Furthermore, in a recent experiment on perceptual awareness of near-threshold tones, pupil dilation exhibited an inverted-U shaped relationship with detection rate, whereby detection was optimal at intermediate pupil sizes (Doll et al., 2024). This pattern aligns with the Yerkes-Dodson law, which posits that moderate arousal levels optimise performance (Yerkes & Dodson, 1908). Importantly, we and others (Eberhardt et al., 2022; Mathôt & Ivanov, 2019) have observed a relationship between pupil size and detection rate and/or accuracy that has a linear component, whereby larger pupils are associated with better detection performance; specifically, referring to Fig. 3c, in our data the relationship between pupil size and sensitivity resembles a combination of an inverted U shape, which may be driven by arousal, and a linear component, which we believe is driven by the optical effect resulting from an increase in pupil size.

In contrast with our results on sensitivity, we found that the relationship between pupil size and criterion (i.e., whether the stimulus was reported as being present) was mediated by power in the beta frequency band. Beta-band oscillations are often associated with motor activity (Engel & Fries, 2010; Jenkinson & Brown, 2011). In our analyses, the criterion is derived from the trial-level responses (a lower criterion is associated with more stimulus-present responses). Since the response here was a button press, it is plausible that the observed mediation is related to motor preparation. Some studies have also linked higher pre-stimulus beta phase coherence to enhanced visual perception (Hanslmayr et al., 2007). However, here we did not observe any relationship between beta and visual sensitivity, further suggesting that the link between beta and criterion may reflect response-related motor activity.

Finally, we have shown positive correlations between pupil size and power in both the alpha and beta bands, replicating findings from previous research (Ceh et al., 2020; Hong et al., 2014; Mathôt et al., 2023; Montefusco-Siegmund et al., 2022; Pilipenko & Samaha, 2024). Alpha power has been more frequently studied, likely because both alpha power and pupil size are linked to similar processes such as visual perception, attention, and arousal. However, our results show a stronger correlation between pupil size and beta power, in line with observations by Mathôt et al. (2023). Considering the known association of beta power with motor activity (Engel & Fries, 2010; Jenkinson & Brown, 2011), one possibility is that this relationship is a reflection of oculomotor control of the iris dilator muscle. Higher beta power has been associated with more tonic contractions of the muscles (Jenkinson & Brown, 2011), which in the iris dilator muscle manifests as a dilation of the pupil. While the exact implications of this relationship remain unclear, its replicability offers a promising direction for further research into the interplay between pupil size and EEG activity patterns.

In conclusion, here we have shown that larger pre-stimulus pupil size is associated with enhanced sensitivity in near-threshold detection. Importantly, this relationship does not appear to be mediated by alpha, beta, or theta power, suggesting an independent influence of pupil size on visual detection. Moreover, we found that pre-stimulus theta suppression is linked to improved detection, which likely mirrors mechanisms typically observed in the alpha-band. Additionally, we observed positive correlations between pupil size and power in both the alpha and beta bands. Our results highlight pupil size changes as a crucial yet still poorly understood factor in visual processing.

